# Network-based atrophy modelling in the common epilepsies: a worldwide ENIGMA study

**DOI:** 10.1101/2020.05.04.076836

**Authors:** Sara Larivière, Raúl Rodríguez-Cruces, Jessica Royer, Maria Eugenia Caligiuri, Antonio Gambardella, Luis Concha, Simon S. Keller, Fernando Cendes, Clarissa Yasuda, Leonardo Bonilha, Ezequiel Gleichgerrcht, Niels K. Focke, Martin Domin, Felix von Podewills, Soenke Langner, Christian Rummel, Roland Wiest, Pascal Martin, Raviteja Kotikalapudi, Terence J. O’Brien, Benjamin Sinclair, Lucy Vivash, Patricia M. Desmond, Saud Alhusaini, Colin P. Doherty, Gianpiero L. Cavalleri, Norman Delanty, Reetta Kälviäinen, Graeme D. Jackson, Magdalena Kowalczyk, Mario Mascalchi, Mira Semmelroch, Rhys H. Thomas, Hamid Soltanian-Zadeh, Esmaeil Davoodi-Bojd, Junsong Zhang, Matteo Lenge, Renzo Guerrini, Emanuele Bartolini, Khalid Hamandi, Sonya Foley, Bernd Weber, Chantal Depondt, Julie Absil, Sarah J. A. Carr, Eugenio Abela, Mark P. Richardson, Orrin Devinsky, Mariasavina Severino, Pasquale Striano, Domenico Tortora, Sean N. Hatton, Sjoerd B. Vos, John S. Duncan, Christopher D. Whelan, Paul M. Thompson, Sanjay M. Sisodiya, Andrea Bernasconi, Angelo Labate, Carrie R. McDonald, Neda Bernasconi, Boris C. Bernhardt

## Abstract

Epilepsy is increasingly conceptualized as a network disorder. In this cross-sectional mega-analysis, we integrated neuroimaging and connectome analysis to identify network associations with atrophy patterns in 1,021 adults with epilepsy compared to 1,564 healthy controls from 19 international sites. In temporal lobe epilepsy, areas of atrophy co-localized with highly interconnected cortical hub regions, whereas idiopathic generalized epilepsy showed preferential subcortical hub involvement. These morphological abnormalities were anchored to the connectivity profiles of distinct disease epicenters, pointing to temporo-limbic cortices in temporal lobe epilepsy and fronto-central cortices in idiopathic generalized epilepsy. Indices of progressive atrophy further revealed a strong influence of connectome architecture on disease progression in temporal lobe, but not idiopathic generalized, epilepsy. Our findings were reproduced across individual sites and single patients, and were robust across different analytical methods. Through worldwide collaboration in ENIGMA-Epilepsy, we provided novel insights into the macroscale features that shape the pathophysiology of common epilepsies.

## Introduction

Perpetual interactions among neuronal populations through the scaffold of axonal pathways promote interregional communication and shape the brain’s structural and functional network organization (Avena-Koenigsberger et al., 2017). This architecture facilitates efficient communication within the brain, and may therefore be profoundly affected by pathological perturbations (Fornito et al., 2015; Stam, 2014). Adopting network science can advance understanding of widespread pathophysiological effects in prevalent disorders and improve diagnostics and prognostication.

The application of neuroimaging to study common epilepsy syndromes has paradigmatically shifted from a focus on individual regions to approaches highlighting network effects, exemplified by classically defined focal epilepsies, such as temporal lobe epilepsy (TLE; Bernhardt et al., 2015; Engel et al., 2013; Gleichgerrcht et al., 2015; Richardson, 2012; Rodriguez-Cruces et al., 2020; Tavakol et al., 2019). While initial work focused on the mesiotemporal lobe, histopathological and neuroimaging studies increasingly detail morphological, structural, and functional compromise beyond this region (Bernhardt et al., 2016; Blanc et al., 2011; Bonilha et al., 2015; Concha et al., 2012; Keller and Roberts, 2008; Labate et al., 2010; Labate et al., 2008; McDonald et al., 2008; Rodriguez-Cruces et al., 2018; Sinjab et al., 2013), which becomes progressively more severe in patients with a longer disease duration (Bernhardt et al., 2009b; Caciagli et al., 2017; Coan et al., 2009; Galovic et al., 2019). Conversely, idiopathic generalized epilepsy (IGE), also known as genetic generalized epilepsy (Scheffer et al., 2017), has been increasingly linked to subtle degrees of structural compromise, mainly in subcortico-cortical circuits (Bernhardt et al., 2009a; Caciagli et al., 2019; Nuyts et al., 2017; O’Muircheartaigh et al., 2012; Wandschneider et al., 2019; Wang et al., 2019). Support for a network perspective also comes from both experimental studies in animal models and electro-clinical observations in patients showing bursts of spike and slow-wave discharges occurring simultaneously over subcortical and cortical areas (Avoli and Gloor, 1982; Blumenfeld, 2003; Gotman et al., 2005). Complementing these observations, basal ganglia atrophy as well as functional changes of the caudate and putamen have been previously noted but warrant further investigation to solidify the understanding of network disruptions in IGE (Bartolini et al., 2014; Keller et al., 2011; Moeller et al., 2008; Paz et al., 2005).

Initiatives such as the Human Connectome Project (HCP; Van Essen et al., 2012) provide normative structural and functional connectivity information from a large sample of healthy individuals. Studying network underpinnings of morphological abnormalities may elucidate brain-wide mechanisms in focal and generalized epilepsies. Hub regions (*i.e.*, nodes with many connections) are a cardinal feature of brain networks and serve as relays to efficiently process and integrate information (Bullmore and Sporns, 2009; van den Heuvel and Sporns, 2011, 2013; Zuo et al., 2012). Their high centrality, however, also makes them vulnerable to pathological processes—the so-called *nodal stress* hypothesis (Crossley et al., 2014; Fornito et al., 2015). Indeed, neurodegenerative and psychiatric research has demonstrated that hubs typically show greater atrophy than locally-connected peripheral nodes. This increased susceptibility to structural damage likely stems from their high metabolic activity and their association with multiple brain networks (Avena-Koenigsberger et al., 2017; Buckner et al., 2009; Crossley et al., 2014; Zhou et al., 2012). We anticipate that models of regional susceptibility can yield significant advances toward our understanding of how connectome architecture configures, to some extent, grey matter atrophy in the common epilepsies.

Complementing the nodal stress hypothesis, in which patterns of cortical atrophy and hub regions appear spatially concomitant, *disease epicenter mapping* can identify one or more specific regions—or epicenters—whose connectivity profile may play a central role in the whole-brain manifestation of focal and generalized epilepsies (Filippi et al., 2020; Raj et al., 2012; Shafiei et al., 2019; Zeighami et al., 2015; Zhou et al., 2012). Among common epilepsies, application of these models to TLE and IGE is justified as both syndromes have been associated with pathophysiological anomalies in mesiotemporal and subcortico-cortical networks and represent conceptual extremes of a focal to generalized continuum of epilepsy subtypes (Bernhardt et al., 2013; Bernhardt et al., 2009a; Keller et al., 2014; O’Muircheartaigh et al., 2012; Weng et al., 2020). Disease epicenter mapping in TLE and IGE may therefore identify syndrome-specific network-level substrates and provide novel insights into how epilepsy-related atrophy patterns are anchored to the connectivity of specific structural and functional subnetworks.

The current work tested the hypothesis that brain network architecture governs whole-brain atrophy in temporal lobe and idiopathic generalized epilepsies. Cortical and subcortical grey matter atrophy patterns were mapped across 19 international sites via ENIGMA-Epilepsy. We also leveraged the HCP dataset to derive high-resolution structural and functional normative brain networks. Two classes of network-based models tested whether, and how, healthy connectome architecture can predict regional susceptibility in the common epilepsies. Our evaluations included: (*i*) nodal stress models, which assessed whether there is a selective vulnerability of hub regions that parallels syndrome-specific atrophy patterns and (*ii*) disease epicenter mapping, which explored the influence of every brain region’s connectivity profile on the spatial distribution of grey matter atrophy in TLE and IGE. In both cases, we model fit was assessed against null models with similar spatial autocorrelation (Alexander-Bloch et al., 2018). To demonstrate clinical relevance, we investigated whether these network-level features could predict spatial patterns of disease progression, inferred here from cross-sectional analysis of disease duration and age interaction effects. We also formulated a patient-tailored adaptation of our network-based models to examine whether network-derived spatial predictors were translatable to individual patients.

## Results

### Data samples

We studied 1,021 adult epilepsy patients (440 males, mean age±SD=36.72±11.07 years) and 1,564 healthy controls (695 males, mean age±SD=33.13±10.45 years) from 19 centres in the international Epilepsy Working Group of ENIGMA (Whelan et al., 2018). Our main analyses focused on two patient subcohorts with site-matched healthy controls: TLE with neuroradiological evidence of hippocampal sclerosis (*n*_HC/TLE_=1,418/732, 341 right-sided focus) and IGE (*n*_HC/IGE_=1,075/289). Details on subject inclusion and case-control subcohorts are provided in the **Methods** and Table 1. Site-specific demographic information is provided in **Table S1**. All participants were aged between 18-70 years.

**Table 1.** ENIGMA Epilepsy Working Group demographics. Demographic breakdown of patient-specific subcohorts with site-matched controls, including age (in years), age at onset of epilepsy (in years), sex, side of seizure focus (TLE patients only), and mean duration of illness (in years). Healthy controls from sites that did not have TLE (or IGE) patients were excluded from analyses comparing TLE (or IGE) to controls.

| **Case-control subcohorts** | **Age (mean±SD)** | **Age at onset (mean±SD)** | **Sex (male/female)** | **Side of focus (L/R)** | **Duration of illness (mean±SD)** |
| --- | --- | --- | --- | --- | --- |
| TLE<br>(*n*=732) | 38.56±10.61 | 16.07±12.27* | 329/403 | 391/341 | 22.74±14.06<sup>a</sup> |
| HC<br>(*n*=1,418) | 33.76±10.54 | — | 643/775 | — | — |
| IGE<br>(*n*=289) | 32.06±10.85 | 16.84±11.25* | 111/178 | — | 15.09±11.70<sup>a</sup> |
| HC<br>(*n*=1,075) | 31.41±9.59 | — | 454/621 | — | — |

<sup>a</sup>Information available in 695/732 TLE patients and 250/289 IGE patients.

### Cortical and subcortical atrophy in the common epilepsies

While the original ENIGMA-Epilepsy study performed a meta-analysis of statistical results submitted by the individual sites, the current study directly analyzed cortical surface and subcortical volume data in all patients and controls. Cortical thickness was measured across 68 grey matter brain regions and volumetric measures were obtained from 12 subcortical grey matter regions and bilateral hippocampi based on the Desikan-Killiany anatomical atlas (Desikan et al., 2006). Surface-based linear models compared atrophy profiles in patients relative to controls; correcting for multiple comparisons using the false discovery rate (FDR) procedure (Benjamini and Hochberg, 1995).

Mirroring ENIGMA-Epilepsy’s meta-analysis of summary statistics comparing neurologically healthy controls to patients with epilepsy, our mega-analysis also revealed widespread cortico-subcortical atrophy patterns in temporal lobe and idiopathic generalized epilepsy syndromes. Specifically, patients with TLE showed profound atrophy in bilateral precentral (*p*_FDR_<2×10^−32^), paracentral (*p*_FDR_<4×10^−30^), and superior temporal (*p*_FDR_<5×10^−21^) cortices as well as ipsilateral hippocampus (*p*_FDR_<2×10^−199^) and thalamus (*p*_FDR_<5×10^−64^, Figure 1A). In contrast, patients with IGE showed atrophy predominantly in bilateral precentral cortices (*p*_FDR_<9×10^−10^) and the thalamus (*p*_FDR_<3×10^−6^, Figure 1B).

**Figure 1.**
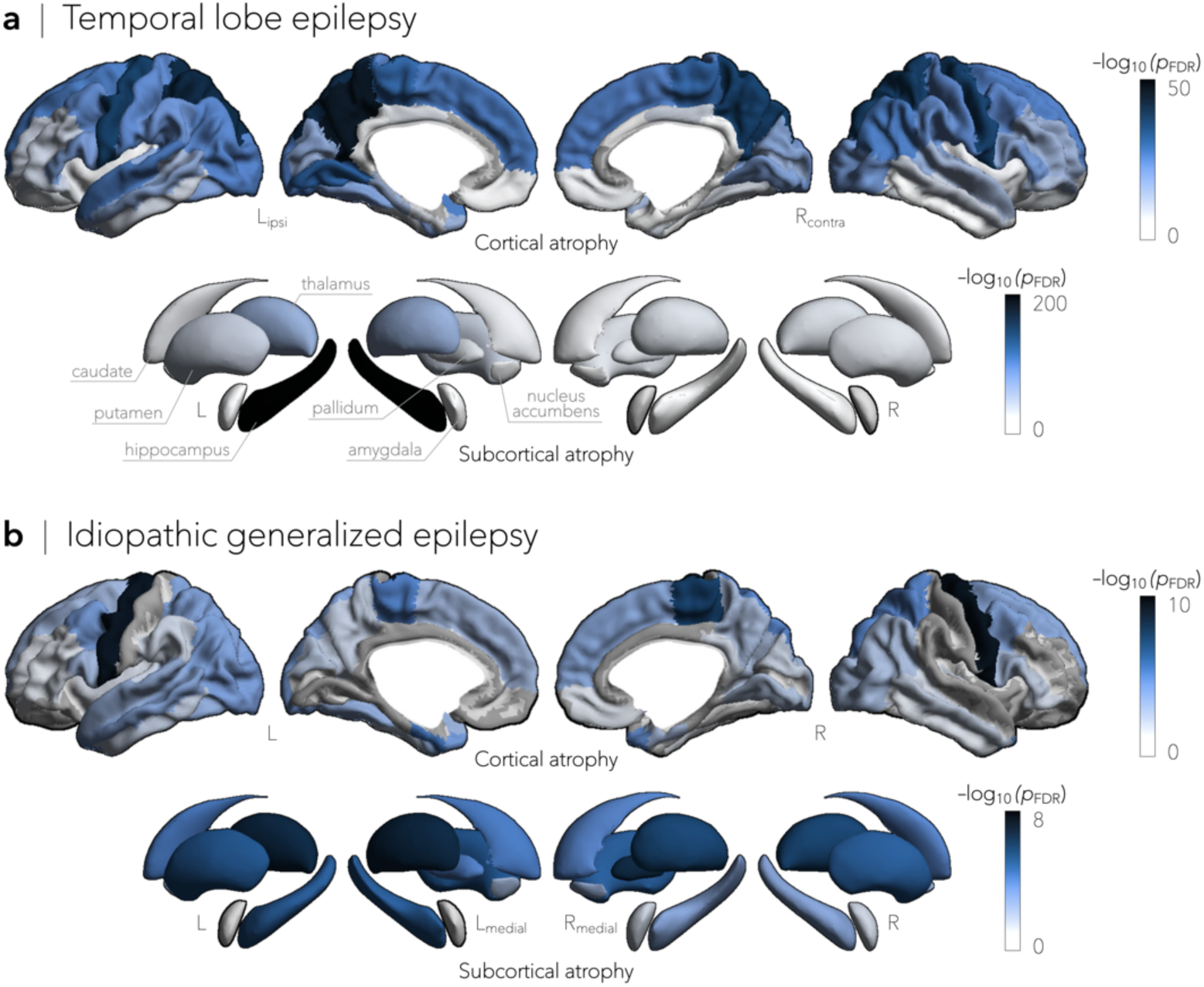
Cortical thickness and subcortical volume in temporal lobe and idiopathic generalized epilepsies. **a** | Cortical thickness and subcortical volume reductions in temporal lobe epilepsy (TLE, *n*=732), compared to healthy controls (*n*=1,418), spanned bilateral precentral (*p*_FDR_<2×10^−32^), paracentral (*p*_FDR_<4×10^−30^), and superior temporal (*p*_FDR_<5×10^−21^) cortices, as well as and ipsilateral hippocampus (*p*_FDR_<2×10^−199^) and thalamus (*p*_FDR_<5×10^−64^). **b** | In contrast, grey matter cortical and subcortical atrophy in idiopathic generalized epilepsy (IGE, *n*=289), relative to controls (*n*=1,075) was more subtle and affected predominantly bilateral precentral cortical regions (*p*_FDR_<9×10^−10^) and the thalamus (*p*_FDR_<3×10^−6^). Negative log10-transformed false discovery rate-corrected (FDR) *p*-values are shown.

### Nodal stress models predict regional susceptibility

Having established patterns of atrophy in TLE and IGE, we evaluated whether these abnormalities were associated with normative network organization. To this end, we obtained high-resolution structural (derived from diffusion-weighted tractography) and functional (derived from resting-state fMRI) connectivity data from a cohort of unrelated healthy adults from the HCP dataset (Van Essen et al., 2012). Details on subject inclusion and matrix generation are provided in the **Methods**.

Echoing established network centrality maps in healthy individuals (van den Heuvel and Sporns, 2013; Zuo et al., 2012), hub regions in the HCP dataset predominated in medial prefrontal, superior parietal, and angular regions (Figure 2A). Nodal stress models, in which spatial similarity between syndrome-related atrophy patterns and degree centrality was compared through correlation analysis (and statistically assessed via non-parametric spin permutation tests, see **Methods**), revealed that cortical thinning in TLE implicated functional (*r*=0.71, *p*_spin_<0.0001) and structural (*r*=0.26, *p*_spin_=0.08) cortico-cortical hubs more strongly than non-hub regions (Figure 2B). In contrast, in IGE, stronger relationships were observed between subcortical volume decreases and functional (*r*=0.50, *p*_spin_=0.06) and structural (*r*=0.68, *p*_spin_<0.01) subcortico-cortical hubs (Figure 2B). To verify stability, we repeated the above correlations across several graph-based nodal metrics (including pagerank centrality and eigenvector centrality) and captured virtually identical associations between atrophy patterns and network centrality measures (**Figures S1, S2**).

**Figure 2.**
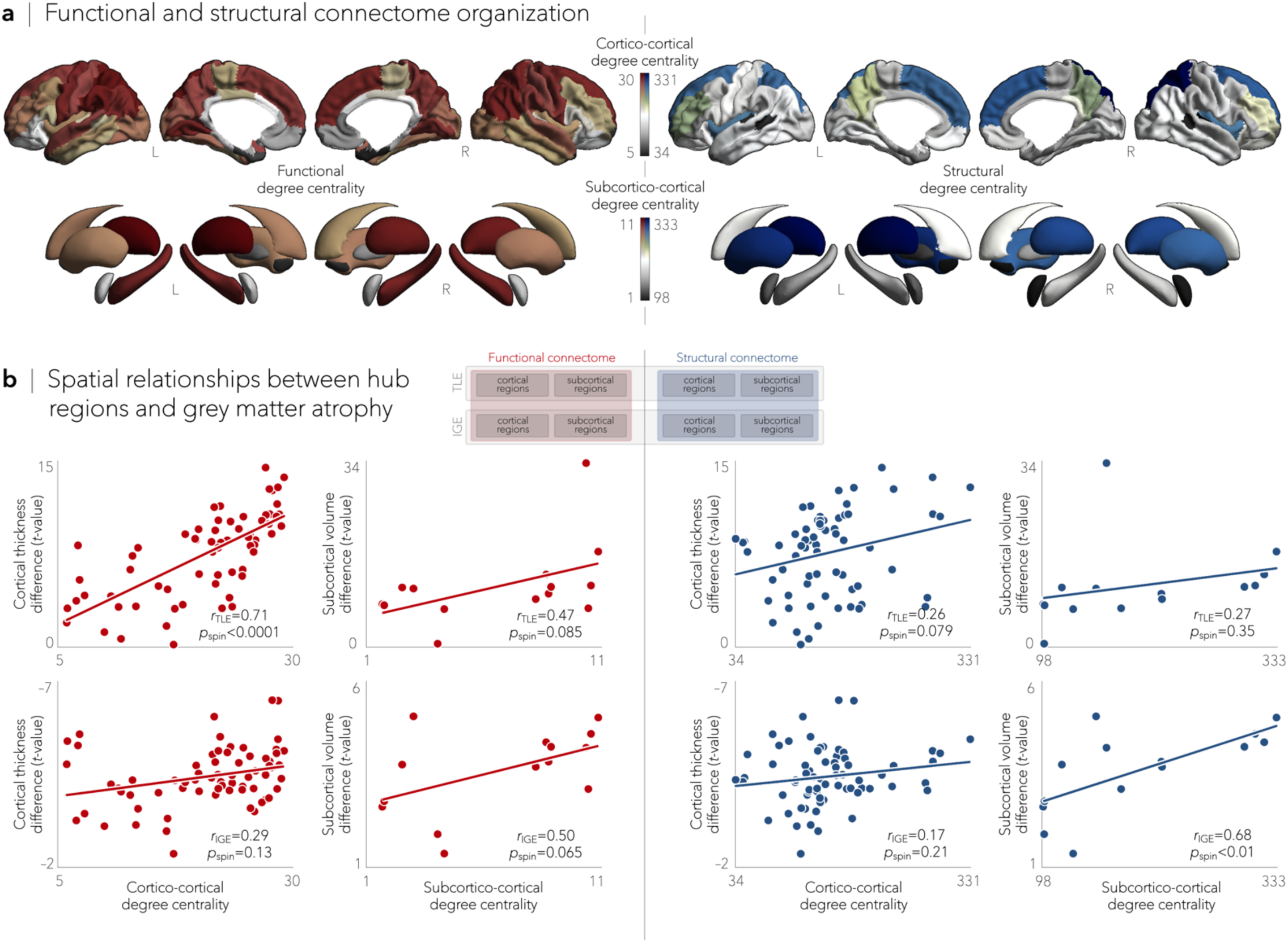
Epilepsy-related atrophy relates to hub organization. **a** | Normative functional and structural network organization, derived from the HCP dataset, was used to identify hubs (*i.e.*, regions with greater degree centrality). **b** | A schematic of the figure layout is pictured in the middle. Grey matter atrophy related to node-level functional (*left*) and structural (*right*) maps of degree centrality, with greater atrophy in hub compared to non-hub regions. Stratifying findings across temporal lobe and idiopathic generalized epilepsies, we observed stronger associations between cortico-cortical functional hubs and cortical atrophy patterns in TLE (*p*_spin_<0.0001) and between subcortical volume loss and subcortico-cortical structural hubs in IGE (*p*_spin_<0.01).

### TLE and IGE have distinct disease epicenters

Since hub regions are more susceptible to atrophy than non-hub regions, we next investigated whether these morphological abnormalities were anchored to the connectivity profile of one, or more, brain regions. To detect syndrome-specific disease epicenters, we systematically compared every region’s functional and structural connectivity profiles to whole-brain patterns of atrophy in TLE and IGE and assessed significance of rankings using spin permutation tests. Cortical and subcortical regions were ranked in descending order based on their correlation coefficients, with highly ranked—and statistically significant—regions being identified as disease epicenters (Figure 3A).

**Figure 3.**
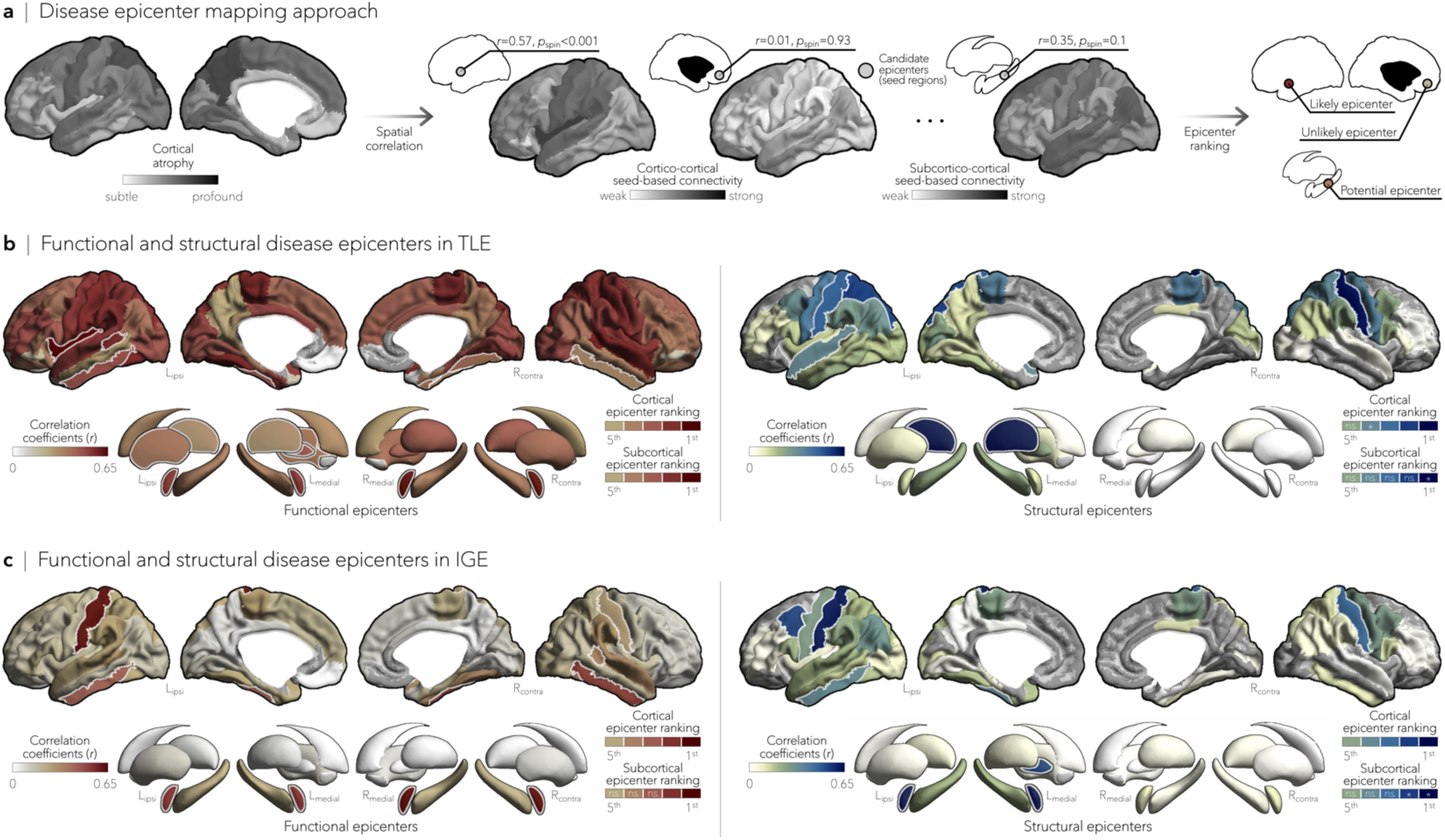
Syndrome-specific disease epicenters. **a** | Disease epicenter mapping schema. Spatial correlations between cortical atrophy patterns and seed-based cortico- and subcortico-cortical connectivity were used to identify disease epicenters in TLE and IGE. Epicenters are regions whose connectivity profiles significantly correlated with the syndrome-specific atrophy map; statistical significance was assessed using spin permutation tests. This procedure was repeated systematically to assess the epicenter value of every cortical and subcortical region, as well as in both functional and structural connectivity matrices. **b** and **c** | Correlation coefficients indexing spatial similarity between TLE- and IGE-specific atrophy and seed-based functional (*left*) and structural (*right*) connectivity measures for every cortical and subcortical region. Regions with significant associations were ranked in descending order based on their correlation coefficients, with the first five regions identified as disease epicenters (white outline). In TLE, ipsilateral temporo-limbic cortices (functional: *p*_spin_<0.05, structural: *p*_spin_<0.1) and subcortical areas, including ipsilateral amygdala (functional: *p*_spin_<0.05), thalamus (functional: *p*_spin_<0.05, structural: *p*_spin_<0.01), pallidum (functional: *p*_spin_<0.05), putamen (functional: *p*_spin_<0.05), and hippocampus (functional: *p*_spin_<0.1), emerged as disease epicenters. In IGE, highest ranked disease epicenters were located in bilateral fronto-central cortices, including postcentral gyri (functional: *p*_spin_<0.05, structural: *p*_spin_<0.05), left (functional: *p*_spin_<0.005, structural: *p*_spin_<0.1) and right amygdala (functional: *p*_spin_<0.005), and left pallidum (structural: *p*_spin_<0.1). ^*^=*p*_spin_<0.1, *n.s.*=non-significant.

In TLE, spatial correlations between atrophy maps and seed-based functional and structural connectivity profiles implicated ipsilateral temporo-limbic cortices (*p*_spin_<0.05) and several ipsilateral subcortical regions as disease epicenters (*p*_spin_<0.05, Figure 3B). Conversely, bilateral fronto-central cortices and the amygdala emerged as epicenters in IGE (*p*_spin_<0.05, Figure 3C). Notably, highest ranked functional and structural disease epicenters in TLE and IGE were significantly connected to hub regions (range of spatial correlations between epicenter-based connectivity and maps of degree centrality: *r*functional=0.68–0.77, *p*_spin_<0.0001; *r*structural=0.27–0.37, *p*_spin_<0.05).

### Cross-sectional indices of disease progression

We adapted the above-described nodal stress and epicenter mapping models to disentangle network factors that contribute to cross-sectional indices of disease progression. Using linear models, we examined effects of age and disease duration on morphological measures within each patient subcohort, paralleling approaches from earlier work (Caciagli et al., 2017).

Patients with TLE showed a negative effect of aging on cortical thickness primarily in bilateral temporo-parietal (*p*_FDR_<0.005) and sensorimotor cortices (*p*_FDR_<0.01), and on subcortical volume mainly in ipsilateral hippocampus (*p*_FDR_<5×10^−7^) and bilateral thalamus (*p*_FDR_<0.05, Figure 4A). Comparing these patterns to structural and functional degree centrality and epicenter maps, strongest correlations were seen with functional cortico- and subcortico-cortical hubs (*r*s=0.64– 0.65, *p*_spin_<0.05), as well as with functional and structural cortical disease epicenters (*r*s=0.41– 0.67, *p*_spin_<0.05, Figure 4B). The main effect of disease duration on atrophy patterns co-localized with regions emerging from the group × age interaction (*e.g.*, bilateral superior parietal cortex, contralateral postcentral gyrus, and ipsilateral hippocampus and thalamus, *p*_FDR_<0.01, **Figure S3A**). While duration effects also showed tendencies for positive correlations with normative network centrality and epicenter values, effects were not significant (**Figure S3B**).

**Figure 4.**
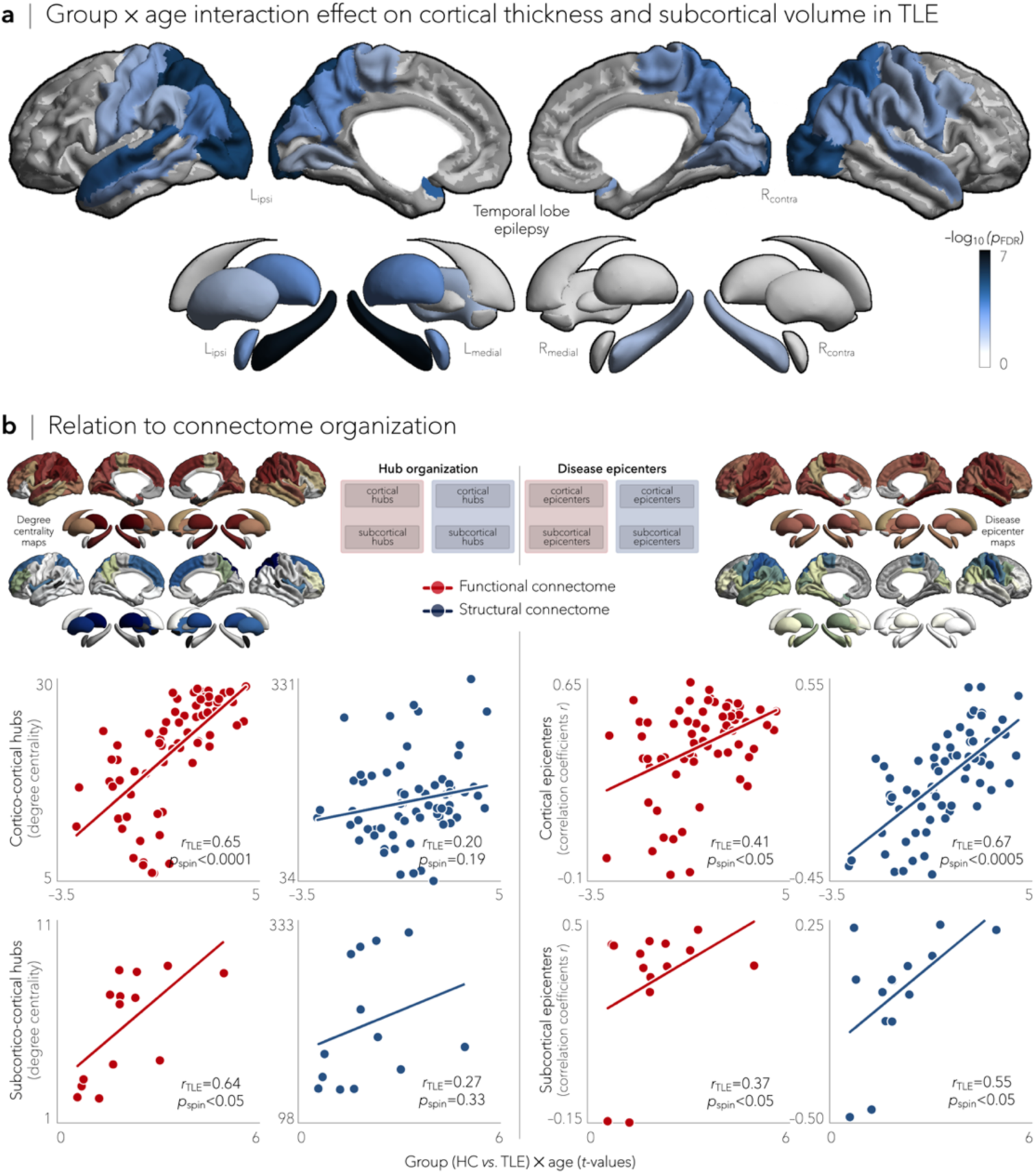
Effects of disease progression on cortical thickness subcortical volume in TLE. **a** | Significant age-related differences on grey matter atrophy between individuals with TLE and healthy controls for all cortical and subcortical regions. Patients with TLE showed a negative effect of aging on cortical thickness in bilateral temporo-parietal (*p*_FDR_<0.005) and sensorimotor (*p*_FDR_<0.01) cortices, and on subcortical volume in ipsilateral hippocampus (*p*_FDR_<5×10^−7^) and bilateral thalamus (*p*_FDR_<0.05). Negative log10-transformed FDR-corrected *p*-values are shown. **b** | A schematic of the figure layout is provided in the middle panel. Scatter plots depict relationships between the age-related effects and functional (*red*) and structural (*blue*) maps of degree centrality (*left*) and disease epicenter (*right*). Significant associations were observed between age-related effects and every hub and epicenter measures, with the exception of structural subcortical degree centrality, suggesting a role of connectome organization on disease progression in TLE.

In IGE, there were no significant effects of aging (**Figure S4A**) or disease duration (**Figure S5A**) on cortical and subcortical atrophy measures, nor of correlations with degree centrality and epicentral maps (**Figures S4B, S5B**). There was a trend towards negative effects of aging and duration of illness on grey matter atrophy, particularly affecting bilateral temporo-parietal and fronto-central cortices (*p*_uncorr_<0.05), as well as right amygdala (*p*_uncorr_<0.1). Exploring the effects of age at onset on morphological abnormalities yielded virtually identical findings in TLE (**Figure S6A**) and IGE (**Figure S6B**), independently.

### Patient-tailored atrophy modelling

We adjusted our network-based models to assess whether normative connectivity organization configures atrophy patterns in individual patients. To do so, we first correlated patient-specific grey matter atrophy maps with normative centrality metrics. We subsequently identified each patient’s structural and functional disease epicenters by keeping brain regions whose healthy connectivity profiles significantly correlated with the patient’s atrophy map.

Although this was expected to result in lower sensitivity due to the increased heterogeneity in atrophy patterns, we observed similar associations between individualized atrophy maps and cortical degree centrality as seen in the group-level analysis. Notably, most stable associations in patients with TLE were seen between cortical hubs and cortical atrophy, whereas most stable relations in patients with IGE were observed between subcortico-cortical hubs and subcortical atrophy (Figure 5A). Findings were robust across both functional and structural connectivity data. Similarly, epicenter results were consistently seen in individual patients, with ipsilateral temporo-limbic and subcortical regions most often emerging as epicenters in TLE, whereas bilateral fronto-central regions, including sensorimotor cortices, were most often identified as epicenters in IGE (Figure 5B).

**Figure 5.**
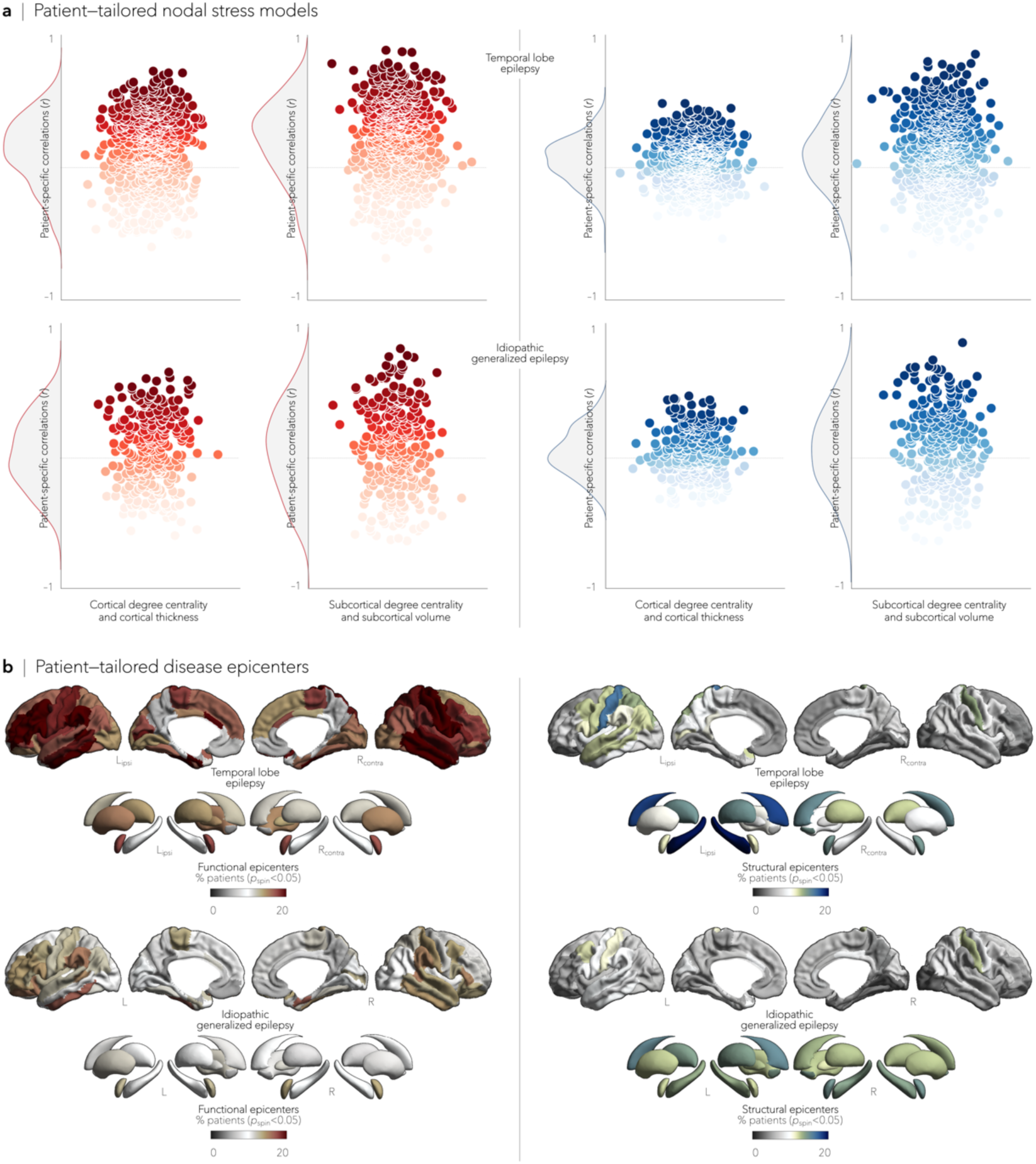
Patient-tailored atrophy modelling. **a** | Patient-specific associations between degree centrality (denoting hub distribution) and individualized atrophy maps showed high stability between cortico- and subcortico-cortical hubs and atrophy in TLE and high stability between subcortico-cortical hubs and subcortical volume reductions in IGE. **b** | We identified patient-specific structural and functional disease epicenters by keeping brain regions whose connectivity profiles significantly correlated with the patient’s atrophy map (*p*_spin_<0.05). In TLE, ipsilateral temporo-limbic regions and subcortical areas (including the hippocampus) were most often identified as epicenters of grey matter atrophy, whereas in IGE, bilateral fronto-central (including sensorimotor cortices) and subcortical areas most often emerged as disease epicenters. Disease epicenters in individual patients strongly resembled those seen across the group as a whole.

### Findings were robust across sites

Syndrome-specific cortical and subcortical atrophy maps were consistent across sites and similar to those obtained from the multi-site aggregation for both TLE (**Figure S7A**) and IGE (**Figure S7B**). Highest stability in TLE was observed for correlations of atrophy with functional cortico- and subcortico-cortical degree centrality (mean±SD: *r*=0.33±0.24 and mean±SD: *r*=0.43±0.26, respectively; **Figure S7C**). Conversely, stability in IGE was highest for correlations with structural subcortico-cortical degree centrality (mean±SD: *r*=0.28±0.22, **Figure S7D**). As observed in the multi-site findings, site-specific ipsilateral temporo-limbic regions and subcortical areas were most often identified as disease epicenters in TLE (**Figure S7E**), whereas bilateral fronto-central cortices, amygdala, and thalamus most frequently emerged as disease epicenters in IGE (**Figure S7F**).

## Discussion

Capitalizing on the largest multi-site epilepsy neuroimaging dataset to date, we tested the hypothesis that grey matter atrophy patterns in temporal lobe and idiopathic generalized epilepsies are related to the brain’s connectome architecture. First, we found that hub regions were overall more susceptible to epilepsy-related atrophy in both TLE and IGE, with cortical hubs most affected in TLE and subcortico-cortical hubs most vulnerable in IGE. These morphological abnormalities were anchored to the connectivity profiles of distinct regions, pointing to temporo-limbic disease epicenters in TLE and fronto-central cortices epicenters in IGE. Assessing markers of disease progression further revealed a dichotomy between TLE and IGE, with a stronger influence of connectome architecture on how the disease unfolds in TLE. Using a patient-tailored adaptation of our network-based models, we confirmed that relationships between atrophy and normative connectivity organization were translatable to individual patients. Findings were highly consistent across sites and methodologies, suggesting robustness and generalizability.

Our study extends prior research by revealing shared and distinct network descriptors of atrophy seen in temporal lobe and idiopathic generalized epilepsies (Whelan et al., 2018). Using a mega-analytic approach, we showed that TLE patients presented with multi-lobar atrophy, affecting fronto-parietal cortices, the hippocampus, and the thalamus, whereas IGE patients presented with more subtle atrophy in precentral regions and the thalamus, despite having ‘normal’ clinical MRIs (Whelan et al., 2018; Woermann et al., 1998). While prior work in histology, electrophysiology, and neuroimaging have provided qualitative descriptions of disease, here we developed network-based models to systematically examine the selective vulnerability of brain regions in the common epilepsies. This approach revealed that epilepsy-related atrophy patterns could be predicted from connectome-derived information, suggesting a vulnerability of cortical hubs to more pronounced atrophy in TLE and an increased susceptibility of subcortical hubs in IGE. This implies that TLE and IGE—despite showing syndrome-specific atrophy patterns—may share common pathophysiological features that result greater atrophy severity in hub regions. Irrespective of cortical or subcortical involvement, the vulnerability of hub regions to grey matter atrophy can be partly ascribed to their disproportionate number of connections and diffuse effect on structural and functional networks (Fornito et al., 2015). Thus, a targeted hub attack, as in the case of epilepsy-related atrophy, may fragment and segregate the network, favoring recurrent seizure activity in temporal lobe and idiopathic generalized epilepsies (Albert et al., 2000; Lariviere et al., 2019b; Larivière et al., 2019).

To elucidate how grey matter atrophy targets hub regions in TLE and IGE, we tested the hypothesis that the underlying connectivity of specific regions—or epicenters—constrains syndrome-specific patterns of atrophy. Other neuroimaging studies of psychiatric and neurodegenerative diseases have employed epicenter mapping techniques to track and predict the spread of atrophy in individual patients (Brown et al., 2019) and disease cohorts (Raj et al., 2012; Shafiei et al., 2019; Zhou et al., 2012). Critically, each epilepsy syndrome harbored distinct epicenters and reflected its own pathophysiology: temporo-limbic cortical regions, ipsilateral to the seizure focus, emerged as epicenters of atrophy in TLE, while fronto-central cortices were identified as disease epicenters in IGE. Syndrome-specific epicenters were functionally and structurally connected to hub regions, suggesting differential mechanisms through which atrophy may spread to hub regions in TLE and IGE.

Taking advantage of large patient cohorts with heterogeneous duration of epilepsy and a wide age range, we could infer the effect of duration of illness on grey matter atrophy and separate out epilepsy-related progression from normal aging. In TLE, progressive atrophy affected temporo-parietal and sensorimotor cortices, as well as the ipsilateral hippocampus and thalamus, as has been previously noted (Bernhardt et al., 2009b; Caciagli et al., 2017; Coan et al., 2009; Galovic et al., 2019; Zhang et al., 2017). Our findings suggest that progressive pathophysiological processes follow the spatial distribution of hub and epicentral regions in temporal lobe, but not idiopathic generalized, epilepsy. We thus speculate that connectome architecture exerts a strong influence on the progression and spread of epilepsy-related cortical damage in patients with TLE. Mechanisms underlying progressive atrophy in TLE remain incompletely understood but may relate to a combination of factors, including seizure-related excitotoxicity and interictal epileptic activity (Bernasconi et al., 2003; Bernhardt et al., 2009a), adverse effects from antiepileptic drugs (Pardoe et al., 2013), and psychosocial impairments (Cascino, 2009). Progressive atrophy findings in IGE—and links to network centrality measures and disease epicenters—were less conclusive. More subtle effects on brain structure in IGE may reflect a less severe disease trajectory compared to focal epilepsy syndromes, and may also arise from the intrinsic heterogeneity across generalized syndromes. It is also possible that generalized epileptiform discharges and seizures diffuse the negative consequences of recurrent pathophysiological activity, and that no single neuronal population bears the brunt of this activity.

Several sensitivity analyses suggested that our findings were not affected by differences in scanners or sites, or graph theoretical metrics. Site and scanner effects were mitigated for the most part using ComBat, a post-acquisition statistical batch normalization process employed to harmonize between-site and between-protocol morphological variations (Fortin et al., 2018). Associations between normative network features and morphological abnormalities, as well as the identification of disease epicenters, were also rather consistently observed in each site independently. Repeating our analyses across a range of graph theoretical metrics yielded virtually identical findings, indicating method invariance of our results. While our ‘big data’ effort was predominantly based on group-level analyses, findings were replicated using patient-specific atrophy maps. Further developed, patient-tailored atrophy models may justify optimism in translating insights from large-scale inference to individual patients, with potential to improve personalized prognostics, treatment monitoring, and epilepsy surgery planning.

We wish to emphasize that our work was made possible by open-access consortia such as the ENIGMA Epilepsy Working Group (Whelan et al., 2018) and the HCP (Van Essen et al., 2012). Although HCP provides a benchmark dataset for structural and functional connectomes in healthy adults, connectivity patterns obtained from this dataset may misrepresent the altered networked architecture typically observed in individuals with epilepsy (Caciagli et al., 2014; Morgan et al., 2011; Wang et al., 2019). Exploiting individualized structural and functional connectome information in patients therefore seems to be the logical next step to improve patient-tailored atrophy models, and ongoing as well as future initiatives to share epilepsy connectome data may facilitate this work. Lastly, the cross-sectional nature of these datasets limited the investigation of effects of disease progression to between-subject effects. Whether within-patient changes in cortical and subcortical atrophy are similarly conditioned by network organization remains an exciting open question and awaits further investigation, ideally in prospective and large-scale collaborative follow-up studies across the spectrum of common epilepsies (Caciagli et al., 2017). We hope that our study lays a foundation for future longitudinal studies to model the predictive path of atrophy in newly diagnosed patients, and further results in network-based clinical tools that can help to understand the pathophysiology of common epilepsies.

## Methods

### ENIGMA participants

We studied 1,021 adult with epilepsy (440 males, mean age±SD=36.72±11.07 years) and 1,564 healthy controls (695 males, mean age±SD=33.13±10.45 years) obtained from the Epilepsy Working Group of ENIGMA (Whelan et al., 2018). Epilepsy specialists at each center diagnosed patients according to the seizure and syndrome classifications of the International League Against Epilepsy (ILAE). Inclusion of adults with TLE was based on the combination of the typical electroclinical features (Berg et al., 2010) and MRI findings typically associated with underlying hippocampal sclerosis. Inclusion of adults with IGE was based on the presence of tonic-clonic, absence, or myoclonic seizures with generalized spike-wave discharges on EEG. No patient had a progressive disease (*e.g.*, Rasmussen’s encephalitis), malformations of cortical development, (*e.g.*, tumors), or underwent prior neurosurgery. Local institutional review boards and ethics committees approved each included cohort study, and written informed consent provided according to local requirements.

### Mega-analysis of cortical thickness and subcortical volumetric data

All participants underwent structural T1-weighted MRI scans at each of the 19 participating centers, with scanner descriptions and acquisition protocols detailed elsewhere (Whelan et al., 2018). Images were independently processed by each center using the standard ENIGMA workflow. In brief, models of cortical and subcortical surface morphology were generated with FreeSurfer 5.3.0 (Dale et al., 1999). Based on the Desikan-Killiany anatomical atlas (Desikan et al., 2006), cortical thickness was measured across 68 grey matter brain regions and volumetric measures were obtained from 12 subcortical grey matter regions (bilateral amygdala, caudate, nucleus accumbens, pallidum, putamen, thalamus) as well as bilateral hippocampi. Data were harmonized across scanners and sites, and statistically corrected for age, sex, and intracranial volume using ComBat—a batch-effect correction tool that uses a Bayesian framework to improve the stability of the parameter estimates (Fortin et al., 2018; Johnson et al., 2007).

Cortical thickness and volumetric measures were *z*-scored relative to site-matched pooled controls and sorted into ipsilateral/contralateral to the focus (TLE only; Liu et al., 2016). As in previous work (Bernhardt et al., 2010; Lariviere et al., 2019a), surface-based linear models compared atrophy profiles in patients relative to controls using SurfStat (Worsley et al., 2009), available at http://mica-mni.github.io/surfstat. Findings were corrected for multiple comparisons using the false discovery rate (FDR) procedure (Benjamini and Hochberg, 1995).

### Spatial permutation tests

The intrinsic spatial smoothness in two given brain maps may inflate the significance of their spatial correlation (Alexander-Bloch et al., 2018). We thus assessed statistical significance of these spatial correlations using spin permutation tests. This framework generates null models of overlap between cortical maps by projecting the spatial coordinates of cortical data onto the surface spheres, applying randomly sampled rotations (10,000 repetitions), and reassigning connectivity values (Alexander-Bloch et al., 2018). The original correlation coefficients are then compared against the empirical distribution determined by the ensemble of spatially permuted correlation coefficients. To compare spatial overlap between subcortical maps, we employed a similar approach with the exception that subcortical labels were randomly shuffled as opposed to being projected onto spheres.

### HCP participants and connectivity matrix generation

We selected a group of unrelated healthy adults (*n*=207, 83 males, mean age±SD=28.73±3.73 years, range=22-36 years) from the HCP dataset (Van Essen et al., 2012). As with the ENIGMA-Epilepsy dataset, high-resolution structural and functional data were parcellated according to the Desikan-Killiany atlas (Desikan et al., 2006). Normative functional connectivity matrices were generated by computing pairwise correlations between the time series (obtained from preprocessed resting-state functional MRI data) of all 68 cortical regions, and between all subcortical and cortical regions. Subject-specific connectivity matrices were then *z*-transformed and aggregated across participants to construct a group-average functional connectome. Moreover, whole-brain streamline tractography (obtained from preprocessed diffusion MRI data) were mapped onto the 68 cortical and 14 subcortical (including hippocampus) brain regions to produce subject-specific structural connectivity matrices. The group-average normative structural connectome was defined using a distance-dependent thresholding, which preserved the edge length distribution in individual patients (Betzel et al., 2019), and was log transformed to reduce connectivity strength variance (Fornito et al., 2016). Details about data collection, scanning parameters, and MRI processing are presented elsewhere (Glasser et al., 2013).

### Nodal stress models

Nodal stress models were derived from spatial correlation analyses between cortical and subcortical syndrome-specific atrophy profiles and normative weighted degree centrality maps. Weighted degree centrality was used here to identify structural and functional hub regions by computing the sum of all weighted connections for every region (higher degree centrality denotes a hub region). Because different centrality measures can describe different topological roles in brain networks (Wang et al., 2018), we replicated the spatial similarity analyses across other nodal metrics, including (*i*) pagerank centrality (proportional to the number of steps (or time) spent at each node), and (*ii*) eigenvector centrality (multiple of the sum of adjacent centralities; *i.e.*, takes into account nodes that are connected to other highly central nodes). To avoid bias in choosing an arbitrary threshold and zeroing potentially useful information, these analyses were carried out on unthresholded connectivity matrices.

### Mapping disease epicenters

Syndrome-specific epicenters were identified by spatially correlating every region’s healthy functional and structural connectivity profiles to whole-brain patterns of cortical atrophy in TLE and IGE (*i.e.*, group-level atrophy maps obtained from surface-based linear models comparing these patient cohorts to controls). This approach was repeated systematically across the whole brain, assessing the statistical significance of the spatial similarity of every region’s functional and structural connectivity profiles to syndrome-specific atrophy maps with spatial permutation tests. Cortical and subcortical epicenter regions were then identified if their connectivity profiles significantly correlated with the disease-specific atrophy map; regions with significant associations were ranked in descending order based on their correlation coefficients, with highly ranked regions representing disease epicenters. As with the nodal stress models, we eliminated matrix thresholding to ensure that connectivity to every brain region was accounted for, thus allowing detection of disease epicenters in areas with subthreshold atrophy. A schematic of the cortical and subcortical disease epicenter mapping approach is displayed in Figure 3A.

To relate disease epicenter findings to the nodal stress model, we performed spatial correlations between epicenter-based connectivity profiles (*i.e.*, whole-brain connectivity seeding from the epicenter region) and maps of cortical degree centrality (showing the spatial distribution of hub regions); strong correlations indicated that disease epicenters were functionally and structurally connected to cortical hub regions. This analysis was individually performed on the two highest ranked functional and structural disease epicenters in TLE and IGE.

### Age- and duration-related effects

To study the effects of cross-sectional indices of disease progression on grey matter atrophy profiles, we first built linear models that included a group and age main effect term as well as a group × age interaction term (Bernhardt et al., 2010; Bernhardt et al., 2009b). We then evaluated age-related differences on cortical thickness and subcortical volume between patients and controls by testing the significance of the interaction term. Linear models independently assessed the effects of duration of epilepsy and age of onset on cortical thickness and subcortical volume measurements in each patient cohort. To investigate the relationship between the effects of disease progression and healthy connectome organization, we compared these age- and duration-related effect maps to normative centrality measures and disease epicenter profiles.

### Patient-tailored atrophy modelling

Cortical thickness and subcortical volume data in patients were *z*-scored relative to healthy controls to generate individualized atrophy maps and were subsequently compared to normative network centrality metrics as in the above analysis. Patient-specific atrophy maps were also used to identify each patient’s structural and functional disease epicenters by keeping the cortical and subcortical regions whose normative connectivity profiles significantly correlated with the patient’s atrophy map. Significance testing for patient-specific epicenters were computed with spin permutation tests with 1,000 repetitions.

### Reproducibility and sensitivity analyses

To address reproducibility of our findings across different sites, we repeated the nodal stress models and disease epicenter analyses in each individual site that provided at least 5 participants per diagnostic group (*n*_TLE/HC_=16 sites, *n*_GE/HC_=10 sites).

## Acknowledgements

The authors would like to express their gratitude to the open science initiatives that made this work possible: (*i*) The ENIGMA-Epilepsy consortium and (*ii*) The Human Connectome Project (Principal Investigators: David Van Essen and Kamil Ugurbil; 1U54MH091657) funded by the 16 NIH Institutes and Centers that support the NIH Blueprint for Neuroscience Research; and by the McDonnell Center for Systems Neuroscience at Washington University.

## Author contributions

*Study conceptualization*: S.L., B.C.B., C.R.M., A.L., M.E.C., N.B., A.B., S.M.S.

*Analysis*: S.L., B.C.B.; individual study sites represented by other co-authors provided preprocessed MRI data and clinical specifiers.

*ENIGMA Epilepsy Leadership*: C.D.W., S.M.S., C.R.M., P.M.T.

*Writing*: S.L., B.C.B.; revised and approved by other listed co-authors.

